# A centrifuge-based method for identifying novel genetic traits that affect root-substrate adhesion in *Arabidopsis thaliana*

**DOI:** 10.1101/2020.05.14.095448

**Authors:** Bethany M Eldridge, Emily R Larson, Laura Weldon, Kevin M Smyth, Annabelle N Sellin, Isaac V Chenchiah, Tanniemola B Liverpool, Claire S Grierson

## Abstract

The physical presence of roots and the compounds they release affect the cohesion between roots and their environment. However, the plant traits that are important for these interactions are unknown and most methods that quantify the contributions of these traits are time-intensive and require specialist equipment and complex substrates. Our lab developed an inexpensive, high-throughput phenotyping assay that quantifies root-substrate adhesion in *Arabidopsis thaliana*. We now report that this method has high sensitivity and versatility for identifying different types of traits affecting root-substrate adhesion including root hair morphology, vesicle trafficking pathways and root exudate composition. We describe a practical protocol for conducting this assay and introduce its use in a forward genetic screen to identify novel genes affecting root-substrate interactions. This assay is a powerful tool for identifying and quantifying genetic contributions to cohesion between roots and their environment.

## INTRODUCTION

Plants secrete compounds that help them adapt to their environment, sense and interact with other organisms, and improve water and nutrient uptake (reviewed in Bais et al., 2006; Sasse et al., 2018; Vives-Peris et al., 2020). These compounds (exudates) can vary in composition based on plant species, developmental stage, and environment (Naveed et al., 2017). Recent studies have shown that some compounds have bioadhesive properties that stick plants and soils together and aggregate soil particles, modifying the microenvironment around plant roots (Galloway et al., 2018). However, understanding how plant biology and physiology contribute to plant-soil interactions can be confounded by multiple factors like other organisms living in the soil that also secrete organic metabolites that can contribute to soil cohesion. Soil composition might affect plant exudate composition and vice versa. Also, plant root morphology including root architecture and the presence of root hairs can alter root-soil interactions (Bailey et al., 2002; Burylo et al., 2012; De Baets et al., 2007; Ghestem et al., 2014; Stokes et al., 2009). This complexity has made it difficult to characterise the relevant plant physiology.

We developed a high-throughput assay that identifies and quantifies the contribution of root hair cells to root-substrate cohesion in a less variable environment than the natural rhizosphere (De Baets et al., 2020). The assay uses a centrifuge to apply force to *Arabidopsis thaliana* seedlings grown on the surface of a sterile medium and measures the force required to detach seedlings from that surface, using a Cox proportional hazards regression (Prentice and Kalbfleisch, 2003) to quantify the adherence of candidate Arabidopsis lines relative to wild-type plants at increasing centrifugal force. Previous use of this assay assessed whether presence or absence of root hairs affected adhesion to the agar (De Baets *et al.,* 2020) but did not address subtle aspects of root hair biology. We now wanted to test whether this assay is sensitive enough to identify other types of traits involved in root-substrate adhesion. To validate this extended use of the assay, we asked if it could detect the adhesive effects in (1) mutants with altered root hair shape; (2) vesicle trafficking mutants with no known root hair morphological defects; (3) exudate compositional mutants with no known root hair phenotypes; and (4) a forward genetic screen of a mutant population to identify novel genes that affect root-substrate adhesion.

Here, we show that this method has the sensitivity to identify how differences in root hair morphology and biological functions that do not present a visible root hair phenotype contribute to substrate cohesion. Using the genetic and molecular tools available in Arabidopsis and working in a sterile environment allows hypotheses about plant specific factors of root-substrate interactions to be tested. While plant-soil cohesion is a complex and dynamic interaction, our assay provides a way to efficiently probe root cellular functions that might be masked by confounding variables in a soil-based study system. Results from this assay provide a platform for more comprehensive studies of plant morphology and physiology that affect root-substrate cohesion.

## MATERIALS AND METHODS

### Materials and Equipment

1. 20% [v/v] bleach solution
2. Sterile distilled water
3. Seed for candidate Arabidopsis lines
4. Half strength Murashige and Skoog basal medium
5. 1% [w/v] sucrose
6. 1% [w/v] agar
7. Laminar air hood
8. 90-mm triple vented Petri Plates
9. Parafilm
10. Growth room
11. Swing-out bucket centrifuge
12. Laboratory safety razor blade
13. Analytical balance

### Plant Material and Growth Conditions

Candidate and wild-type lines of the Columbia-0 (Col-0) ecotype of *Arabidopsis thaliana* single-gene mutants were obtained from the Nottingham Arabidopsis Stock Centre (NASC; Nottingham, UK) or were stocks maintained in the lab (**Table 1**). Homozygous lines were used in all experiments.

**Table 1.** *Arabidopsis thaliana* mutant lines used in this study.

| *Mutant type* | *Mutant* | *AGI code<sup>a</sup>* | *Mutant line* | *Referenced in* |
| --- | --- | --- | --- | --- |
| Root hair shape | *cow1-3* | At4g34580 | SALK_002124 | Böhme *et al.* (2004) |
|  | *csld3-1* | At3g03050 | CS899 | Wang *et al.* (2001) |
|  | *lrx1-4* | At1g12040 | SALK_057038 | Baumberger *et al.* (2001) |
|  | *rol1-2* | At1g78570 | CS16373 | Diet *et al.* (2006) |
|  | *gdi1-2* | At3g07880 | SALK_035400 | Kang *et al.* (2017) |
| Vesicle trafficking | *chc1-2* | At3g11130 | SALK_103252 | Kitakura *et al.* (2011) |
|  | *chc2-3* | At3g08530 | SALK_151638 | Mbengue *et al.* (2016) |
|  | *syp121* | At3g11820 | - | Collins *et al.* (2003) |
|  | *syp122-1* | At3g52400 | SALK_008617 | Assaad *et al.* (2004) |
| Root exudate composition | *jin1-9* | At1g32640 | SALK_017005 | Anderson *et al.* (2004) |
|  | *pdr2* | At4g15230 | SAIL_811_F08 | Badri *et al.* (2008) |
|  | *pft1-3* | At1g25540 | SALK_059316 | Kidd *et al.* (2009) |
| Unknown | *abcg43-1* | At4g15236 | N75206<sup>b</sup> | This study |
|  | *abcg43-2* | At4g15236 | SALK_201207 | This study |
|  | *abcg43-3* | At4g15236 | SALKseq_30713 | This study |

For all experiments, seeds were surface sterilised for 15 min in 20% [v/v] bleach and sterile water solution, followed by five washes in sterile water. All sterilised seeds were stratified in the dark at 4°C for 24-48 h in Eppendorf microtubes containing sterile water. All plants were reared in a growth room at 21-22°C with a continuous photoperiod (light intensity = 120-145 μmol m^−2^ s^−1^) at 60% humidity.

Half strength Murashige and Skoog basal medium was supplemented with 1% [w/v] sucrose pH adjusted to 5.7, solidified with 1% [w/v] agar. A 30 ml volume of sterilized media (at ~45 – 50°C) was pipetted into 90-mm triple vented Petri plates and left to set for ~30 min in a laminar air hood. Sterilized seeds were sown onto Petri plates containing the sterile medium and sealed with Parafilm. Plates were then placed in stacks of five at ~80° from the horizontal (lid side up) to encourage vertical growth along the gel surface. Plates were incubated in the growth chamber for 5-6 days before centrifugation. Plants for genomic DNA extractions were grown in compost (three parts Levington F3 compost and one-part J Arthur Bowers horticultural silver sand).

### Centrifuge Assay Set-Up

To measure the adhesion of Arabidopsis seedling roots using a centrifuge, seedling roots must not penetrate the gel or touch each other, as both events could interfere with seedling detachment during centrifugation. Therefore, ten seeds were sown 1-cm apart on each plate in two parallel rows (**Figure 1Ai**). The plates were then grown for 5-6 days (**Figure 1Aii**) and seedlings that were touching each other, had grown into the gel, or not uniform in size were omitted from the assay. The seedlings were then numbered so that plate positional effects could be included as a potential covariate in the final analysis (**Figure 1Bi**). Plates were inverted and placed into a swing-out bucket centrifuge (Beckman Allegra X-30R Centrifuge) that held four plates at a time and subjected to increments of increasing centrifugal force at 720, 1018, 1247, 1440 and 1611 RPM for one minute each (**Figure 1Bii**). The proportion of seedlings that detached from the gel surface was recorded between each centrifugal speed; a detachment event was recorded when a seedling had partially or fully peeled away from the gel (**Figure 1Biii**). If the gel shattered during the assay, seedlings that had not detached from the gel were not recorded and seedlings that remained adhered to the gel after the maximum centrifugal speed were censored in the analysis. During method optimisation we encountered some common methodological challenges for which we provide troubleshooting options in **Table S1**.

**FIGURE 1.**
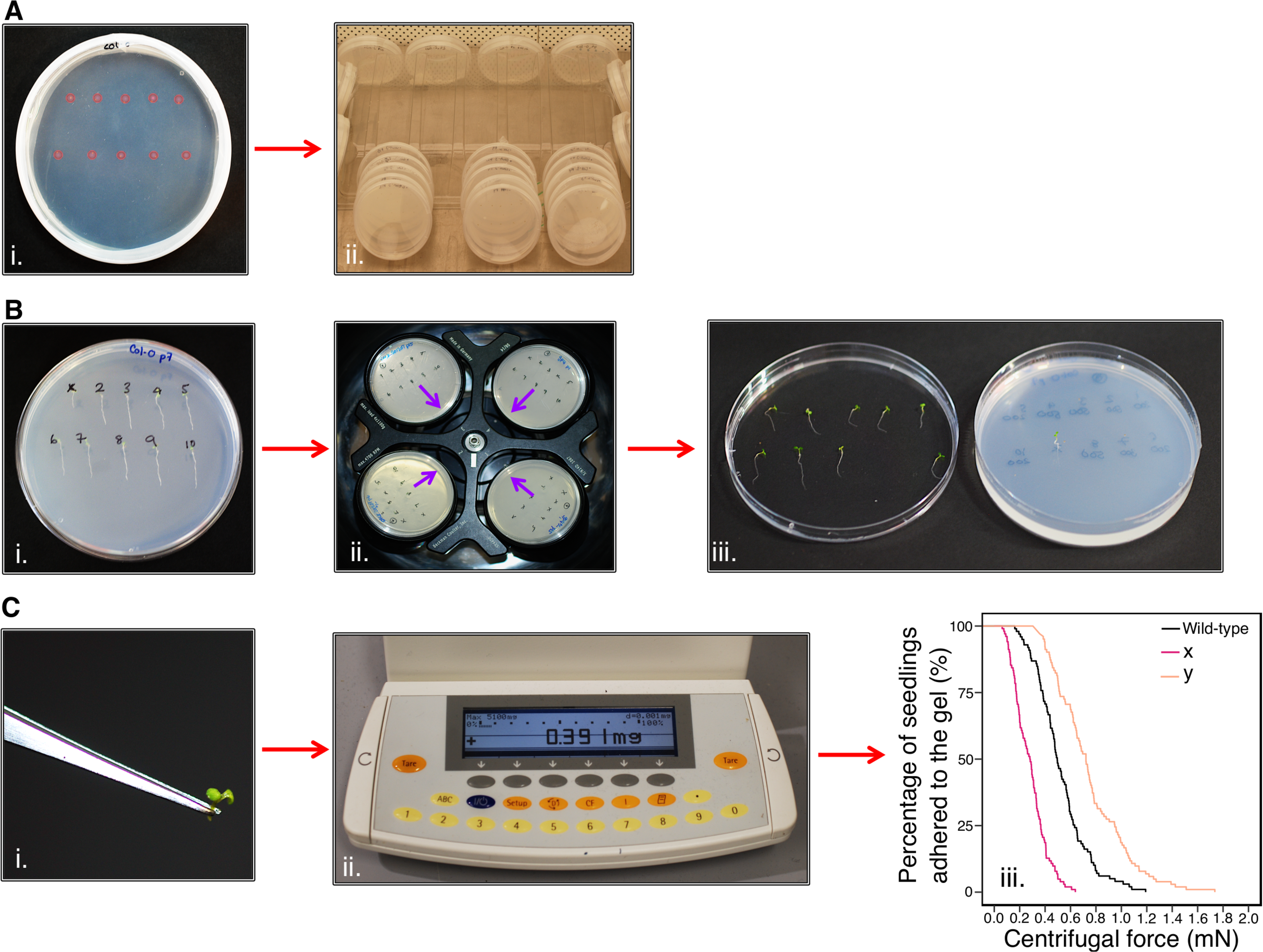
Centrifuge assay workflow. **(A)** i. Ten seeds (highlighted in pink) are sown onto the surface of sterile, solid gel growth medium in a single 90 mm Petri plate in two horizontal rows. ii. The plates are stacked in groups of five and orientated vertically at approximately 80° to encourage the roots to grow down the surface of the gel medium. Plates are grown in a growth chamber with constant light (120-145 μmol m^−2^ s^−1^) conditions at 22°C and 60% relative humidity. **(B)** After 5-6 days, i. seedlings are visually analysed and numbered. ii. Petri plates are placed into a swing-out-bucket centrifuge in an inverted orientation with their roots pointing inward (indicated by the purple arrows). iii. After seedlings have been subjected to each one-minute pulse of increasing centrifugal speed, seedling detachment is recorded. **(C)** i. The aerial tissue mass of each seedling is determined using ii. an analytical scale to iii. determine the root-gel adhesion properties of candidate lines are compared to wild-type to assess whether they have increased (e.g. line y) or decreased (e.g. line x) adhesion to the sterile gel.

### Calculating Seedling Detachment Force

To calculate the centrifugal force acting on a seedling, the aerial tissue mass for each seedling was separated from the root by making an incision at the hypocotyl-root junction using a laboratory safety razor blade and immediately measured using an analytical balance (**Figure 1Ci, ii**). The aerial tissue weight of each seedling was measured because a plant with heavier aerial tissue experiences more force than a lighter one subjected to the same centrifugal speed. Centrifugal force, Fc (mN) acting on a seedling was calculated as previously reported in De Baets et al. (2020). Aerial tissue weight (kg), the angular velocity (*ω*) and the distance between the seedling and the axis of rotation on the centrifuge (radius = 0.07m) were used to give the following equation:

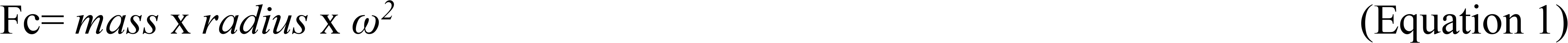

### Identification of *abcg43-1* Mutant T-DNA Insert

Genomic DNA was extracted from a pool of vegetative tissue taken from ten three-week old plants using an adapted protocol (Healey et al., 2014) for Illumina Next-Generation two x paired-end sequencing conducted at the Bristol Genomics Facility (University of Bristol, UK). The Illumina TruSeq Nano LT gDNA kit (Illumina Inc) was used to generate a genomic DNA sequencing library following the manufacturer’s protocol. The final library was diluted to a loading concentration of 1.4 pM for cluster generation and 2 × 150 bp paired-end sequencing on the Illumina NextSeq500 system (Illumina Inc) alongside a 5% PhiX spike-in control library. Read summary statistics were generated using the RTA 2.4.6 Primary Analysis Software (Illumina Inc). The read summaries were analysed in the Sequencing Analysis Viewer (Illumina Inc). Filtered paired reads were subject to paired-end alignment using the Bowtie2 2.3.4.2 aligner (Langmead and Salzberg, 2012). A bespoke reference genome was produced that combined the TAIR10 Arabidopsis genome (Lamesch et al., 2012) and the pROK2 vector sequence (Baulcombe et al., 1986). Alignments were viewed in the Integrative Genomics Viewer IGV 2.3 (Robinson et al., 2011).

For Sanger sequencing, DNA was extracted from a pool of vegetative tissue taken from two-week-old plants using a modified Edwards prep (Edwards et al., 1991). High fidelity PCR was conducted on the *abcg43-1* line using the Q5 High-Fidelity 2X Master Mix (NEB). PCR products were purified and extracted from a 1% agarose gel using the QIAquick Gel Extraction Kit (QIAGEN), following the manufacturer’s instructions. Purified PCR product plus the *abcg43-1* forward, reverse or border genotyping primer (**Table S2**) was used for Sanger sequencing (using the Mix2Seq overnight sequencing kit, Eurofins Genomics). Chromatograms and FASTA files were obtained from Eurofin Genomics and following manual low-quality end trimming, the final sequences were aligned to the pROK2 vector sequence and *ABCG43* gene sequence using the MUSCLE alignment tool (Madeira et al., 2019).

### Genotyping

To genotype the *abcg43* mutant alleles, genomic DNA was extracted from the vegetative tissue of two-week-old plants using a modified Edwards prep (Edwards et al., 1991). T-DNA border and gene-specific primers were used in PCR analyses to confirm the genotype (**Table S2**).

### Morphological Root Trait Analyses

Five-day-old Arabidopsis seedlings were imaged with a Leica MZ FLIII microscope (Leica) with dark-field lighting. Root hair length and density were measured using microscope images of five-day-old Arabidopsis seedlings and Fiji version 1.0 (Schindelin et al., 2012) and the Bio-Formats Importer plugin. At least two experimental repeats were conducted; in each experiment, we imaged eight to ten individual plants from each line and measured at least 30 root hairs per plant.

### Statistical Analyses

All statistical analyses were conducted using RStudio, version 1.1453 (R Core Team, 2014) and all graphs were generated using the R package ggplot2 (Wickham, 2016).

#### Centrifuge assay

Survival analysis was used to study when seedlings detach from the sterile gel medium (i.e. seedling detachment is the event) at increasing force (i.e. force is modelled rather than time). We used the Cox proportional hazards (PH) regression function, coxph, within the survival package in R to statistically test for differences between the detachment of experimental seedlings relative to the wild-type control (Col-0), using centrifugal force as the predictor variable (**Figure 1Ciii)**. We incorporated seedling position and individual plate number as covariates within our Cox PH regression models to account for any environmental heterogeneity such as minor inconsistencies in the water content or gelling of the medium. When these covariates had no significant effect, they were removed from the model. Seedlings that remained adhered to the gel after the maximum centrifugal speed were censored in our Cox PH regression model. Comparisons of different candidate lines relative to the wild type line were analysed *a priori*; therefore, a series of contrasts were set up using the R function contr.treatment rather than using post-hoc testing methods.

For each Cox PH regression model run, we report the P value of the Wald Statistic (*z*-score) and the hazard ratio with the upper and lower bound confidence intervals. Hazard ratios are exponentiated coefficients that give the instantaneous risk of root-gel detachment (Cox & Oakes, 1984; Bradburn *et al.*, 2003; Spruance *et al.*, 2004). For each Cox PH regression model conducted, we used the hazard ratio as a measure of effect size to assess differences between the root-gel adhesion of candidate lines and wild type (Cox & Oakes, 1984). Wild type was used as the baseline group for all models, with a hazard ratio of one. A candidate line with higher risk of detachment from gel than wild type will have a hazard ratio above one. Conversely, lines with a lower risk of detachment than wild type will have a hazard ratio below one.

For all Cox PH regression models, the assumption of proportionality was satisfied and an alpha level of 0.01 was used. The root-gel detachment of at least 70 plants from each line was assessed within a single experiment.

#### Root hair analyses

Two sample t-tests were conducted using the t.test function in R to test for differences in the mean root hair density or mean root hair length (of eight to ten individual plants from each line) between candidate lines relative to wild type (Col-0). In all cases, the assumptions of normality and homoscedasticity were satisfied. Because comparisons of different mutant lines relative to the wild type line were *a priori*, a series of contrasts were conducted using the contr.treatment function in R. To prevent type I errors from multiple testing, we applied the Bonferroni method and adjusted the alpha level to 0.025 (0.05/2).

## RESULTS

### Root Hair Morphological Effects on Root-Substrate Adhesion

We screened mutant lines with known root hair morphology defects to ask if the centrifuge assay could determine if root hair shape affects root-substrate adhesion. The mostly bald mutant *csld3*, which encodes the CESA-like 3D protein (Favery et al., 2001; Wang et al., 2001; Yang et al., 2020) was included as a root hairless example. For lines that produce short or malformed root hairs, we included the *gdi1-2* Rho GTPase GDP dissociation inhibitor mutant, *rol1-2* rhamnose biosynthesis 1 mutant allele, *can of worms 1* (*cow1-3*) phosphatidylinositol transferase protein mutant, and *lrx1-4* leucine rich extensin 1 mutant (Baumberger et al., 2001; Böhme et al., 2004; Carol et al., 2005; Diet et al., 2006; Grierson et al., 1997). The *rol1-2* and *gdi1-2* mutants have short root hairs and the root hairs on *gdi1-2* plants can also branch (Diet et al., 2006; Kang et al., 2017; Parker et al., 2000; Ringli et al., 2008); *cow1-3* plants have short, wide root hairs that can branch; and *lrx1-4* plants can develop root hairs that abort, swell, branch, or collapse in a growth condition-dependent manner (Baumberger et al., 2001; Böhme et al., 2004; Grierson et al., 1997; Parker et al., 2000). Assay results for ectopic and long root hairs were reported previously (De Baets et al., 2020). Under our growth conditions, we confirmed the reported phenotypes for each mutant line (**Figure 2A**). Wild type was used as the baseline group for all centrifuge experiments, with a hazard ratio of one. Therefore, a candidate line with a higher risk of detachment from the gel than wild type will have a hazard ratio above one, while lines with a lower risk of detachment than wild type will have a hazard ratio below one.

**FIGURE 2.**
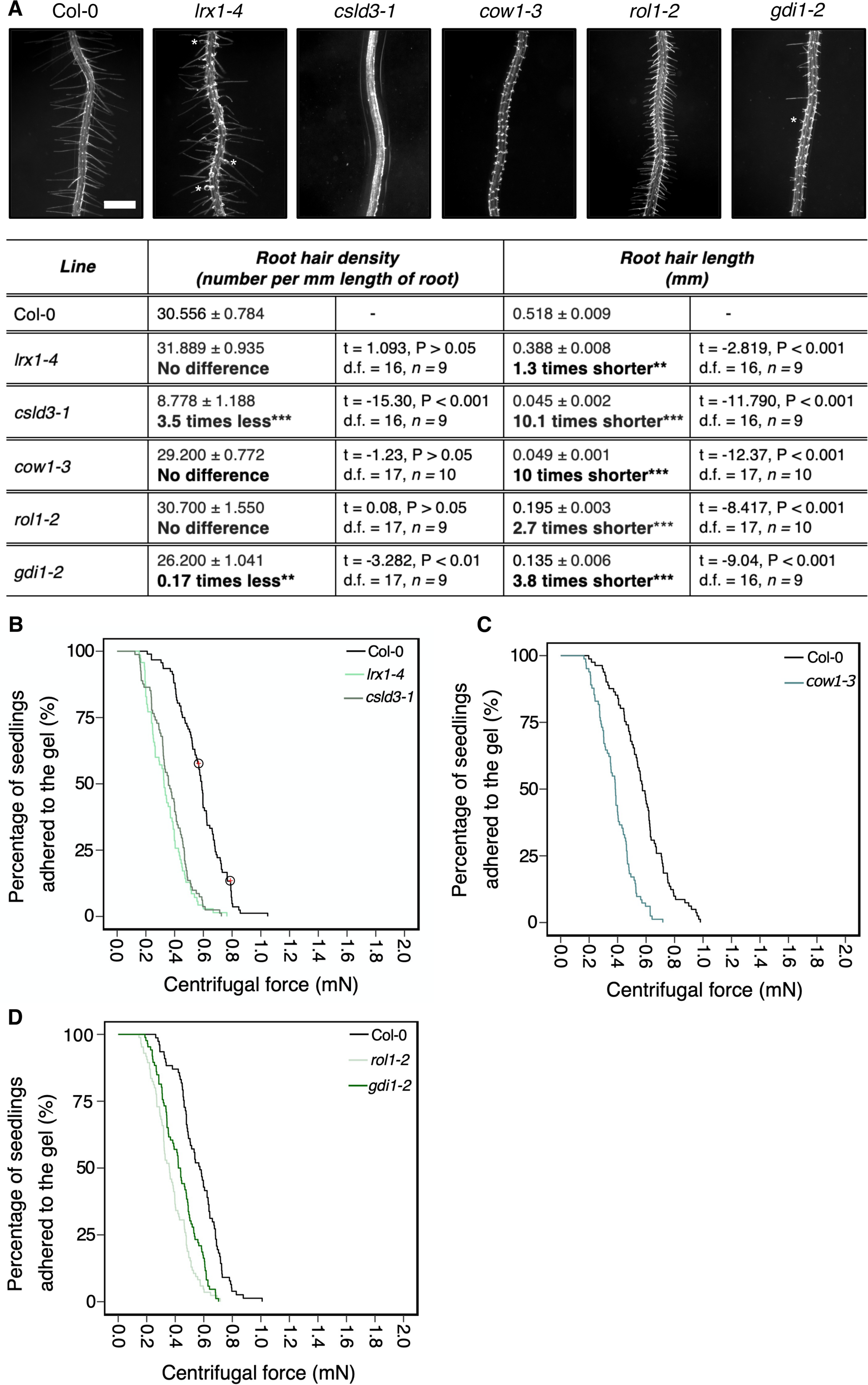
Physical root hair properties contribute to root-substrate adhesion in *Arabidopsis thaliana.* **(A)** Root hair phenotypes of 5-d-old wild-type (Col-0), *lrx 1-4, csld3-1, cow1-3, rol1-2,* and *gdi1-2* seedlings grown on gel medium and statistical comparisons of mean root hair density (number per mm length of root) and mean root hair length (mm) for each mutant line relative to wild type. White asterisks on the root hair images indicate the characteristic root hair bulging phenotype in the *lrx1-4* mutant and the short, branching root hair phenotype in *gdi1-*2. Scale bar = 0.5 mm. In the table, the mean *±* standard error is given for root hair density and root hair length of wild type and each mutant as well as a mean comparison to wild type, which is listed in bold. “No difference” is stated when there was no statistically significant difference between wild type and a mutant line. The statistical output of each univariate linear model is given. Alpha = 0.025, statistical significance codes: ‘***’ = < 0.001, ‘**’ = < 0.01. The results shown are from a representative experiment for at least two independent experiments. Survival curves displaying the proportion of seedlings that adhered to the gel at increasing centrifugal force for **(B)** 92 wild type (Col-0 – black); 81 *lrx1-4* (light green); 70 *csld3-1* (green); **(C)**82 wild type (Col-0 - black); 83 *cow1-3* (green); and **(D)**77 wild type (Col-0-black); 85 *rol1-2* (light green); and 86 *gdi1-2* (dark green). Circled red crosses on the survival curves represent seedlings that remained adhered to the gel after the maximum centrifugal speed (1611 RPM). Each graph shows a representative experiment for at least two independent experiments showing a statistically significant difference in adhesion between mutant lines relative to wild type (Cox PH regression; alpha = 0.01). A single experiment included > 70 biological replicates for each candidate line.

Consistent with previous results (De Baets et al., 2020), bald (*csld3*) seedlings detached at lower centrifugal forces relative to wild-type seedlings, with 4.6 times the risk of detachment than wild-type plants (P < 0.001 – **Table 2**; **Figure 2B**). Mutants with deformed root hair phenotypes including *gdi1-2, rol1-2, cow1-3* and *lrx1-4* had risks of detachment that were 2.5, 4, 3.8 and 3.9 times that of wild-type plants, respectively (for all lines P < 0.001 - **Table 2**; **Figure 2B-D**). These results indicated that the centrifuge assay can quantify the effects of root hair shape on root-substrate cohesion.

**Table 2.** Output of the Cox PH regression models comparing the root-gel detachment of root hair shape, vesicle trafficking and root exudate composition mutants relative to wild type.

| *Mutant type* | *Mutant* | *Wald test (z-score) and corresponding P value* |  | *Hazard ratio (95% CI)* |
| --- | --- | --- | --- | --- |
| Root hair shape | *cow1-3* | *z* = 7.350<br>*P* < 0.001 | *** | 3.790<br>(2.657, 5.407) |
|  | *csld3-1* | *z* = 8.644<br>*P* < 0.001 | *** | 4.636<br>(3.274, 6.564) |
|  | *gdi1-2* | *z* = 5.199<br>*P* < 0.001 | *** | 2.459<br>(1.752, 3.452) |
|  | *lrx1-4* | *z* = 8.049<br>*P* < 0.001 | *** | 3.926<br>(2.814, 5.478) |
|  | *rol1-2* | *z* = 7.927<br>*P* < 0.001 | *** | 4.036<br>(2.858, 5.698) |
| Vesicle trafficking | *chc1-2* | *z* = 8.238<br>*P* < 0.001 | *** | 4.482<br>(3.137, 6.405) |
|  | *chc2-3* | *z* = 7.653<br>*P* < 0.001 | *** | 3.941<br>(2.774, 5.599) |
|  | *syp121* | *z* = 7.702<br>*P* < 0.001 | *** | 3.593<br>(2.595, 4.975) |
|  | *syp122-1* | *z* = 7.199<br>*P* < 0.001 | *** | 3.270<br>(2.369, 4.515) |
| Root exudate composition | *jin1-9* | *z* = 0.737<br>*P* > 0.05 | ns | 1.124<br>(0.824, 1.537) |
|  | *pdr2* | *z* = -5.765<br>*P* < 0.001 | *** | 0.379<br>(0.273, 0.528) |
|  | *pft1-3* | *z* = 3.315<br>*P* < 0.001 | *** | 1.698<br>(1.242, 2.323) |
| Unknown | *abcg43-1* | *z* = -7.823<br>*P* < 0.001 | *** | 0.245<br>(0.172, 0.348) |
|  | *abcg43-2* | *z* = -6.049<br>*P* < 0.001 | *** | 0.355<br>(0.253, 0.496) |
|  | *abcg43-3* | *z* = -7.179<br>*P* < 0.001 | *** | 3.496<br>(0.203, 0.403) |

\*\*\* refers to a statistically significant difference $\leq 0.001$ and **ns** no statistically significant difference $\geq 0.05$ relative to wild type, respectively.

### Vesicle Trafficking Mutations Affect Root-Substrate Adhesion

We hypothesized that Arabidopsis mutants with defective vesicle trafficking pathways might alter cell wall and apoplast characteristics that contribute to root-substrate interactions and adhesion. We chose the secretory Soluble NSF (N-ethylmaleimide sensitive fusion protein) Attachment proteins (SNAP) REceptor (SNARE) mutants *syp121* and *syp122-1*, and the endocytic mutants, *chc1-2* and *chc2-3* as candidates for our assay. SYP121 is the primary secretory SNARE found on the plasma membrane (Assaad et al., 2004; Geelen et al., 2002) and has characterized aboveground phenotypes, including small rosettes and stomatal mobility defects (Eisenach et al., 2012; Larson et al., 2017); however, the mutant does not have a reported root hair phenotype. Although the related SNARE SYP122 is thought to share partial functionality with SYP121, the *syp122-1* mutant does not share these phenotypes with *syp121* and a recent proteomic analysis reported differences in their vesicle cargoes, which could contribute to their functional independence (Waghmare et al., 2018). The *chc1-2* and *chc2-3* mutants are defective in the heavy chain subunits of the clathrin coat complex, which is required for vesicle traffic at the plasma membrane (Kitakura et al., 2011; Mbengue et al., 2016). Endo- and exocytic rates are impaired in both mutants (Larson et al., 2017), but no root hair phenotypes have been reported. For the centrifuge assay, it was important to select mutants that do not have root hair phenotypes given the effect root hair morphology has on root-gel adhesion. Therefore, we evaluated the root hairs of these vesicle trafficking mutants and found that the root hairs of these lines did not significantly differ from wild-type plants (**Figure 3A**). Overall, the risk of detachment for *syp121, syp122-1, chc1-2* and *chc2-3* plants was 3.6, 3.3, 4.5 and 3.9 times that of wild type, respectively (for all lines P < 0.001 – Table 2; **Figure 3B,C**). These results indicated that the centrifuge assay can be used to probe for intracellular molecular machinery that affect root-substrate interactions.

**FIGURE 3.**
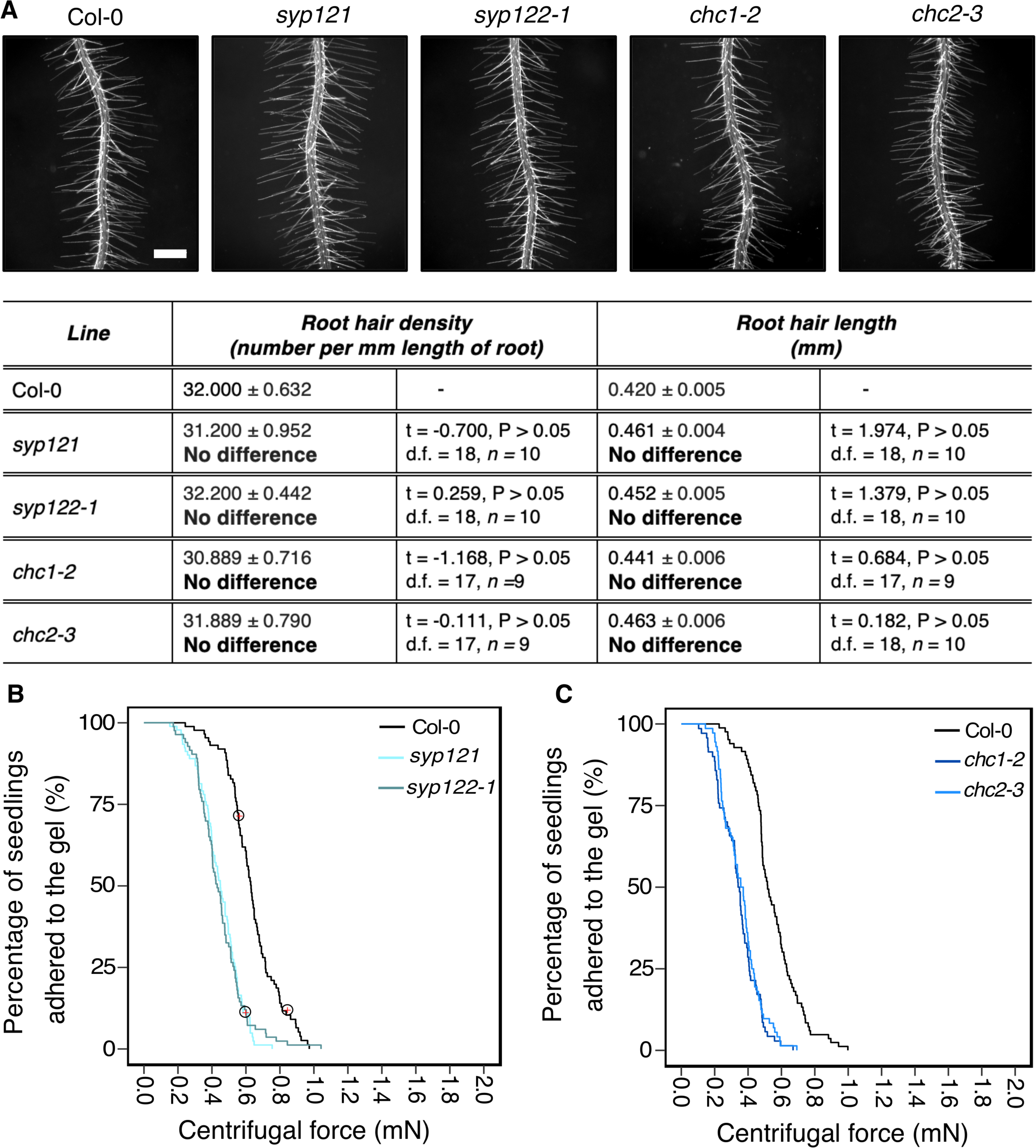
Vesicle trafficking mechanisms that contribute to root-substrate adhesion in *Arabidopsis thaliana.* **(A)** Root hair phenotypes of 5-d-old wild-type (Col-0), *syp121, syp122-1, chc1-2* and *chc2-3* seedlings grown on a gel medium and statistical comparisons of mean root hair density (number per mm length of root) and mean root hair length (mm) for each mutant line relative to wild type. Scale bar = 0.5 mm. In the table, the mean *±* standard error is given for the root hair density and root hair length of wild type and each mutant as well as a mean comparison to wild type, which is listed in bold. “No difference” is stated when there was no statistically significant difference between wild type and a mutant line. The statistical output of each univariate linear model is given. Alpha = 0.025. The results shown are from a representative experiment for at least two independent experiments. Survival curves displaying the proportion of seedlings that adhered to the gel at increasing centrifugal force for **(B)**87 wild type (Col-0 – black); 91 *syp121* (turquoise); 83 *syp122-1* (blue/grey); and **(C)**83 wild type (Col-0 – black); 70 *chc1-2* (dark blue); and 72 *chc2-3* (medium blue). Red crosses circled on the survival curves represent seedlings that remained adhered to the gel after the maximum centrifugal speed (1611 RPM). Each graph shows a representative experiment for at least two independent experiments showing a statistically significant difference in adhesion between all mutant lines relative to wild type (Cox PH regression; alpha = 0.01). A single experiment included > 70 biological replicates for each candidate line.

### Exudate Composition Changes Root Detachment Rates

Over 20% of plant assimilated carbon is released as root exudates (Huang et al., 2016). These are complex mixtures of diverse molecular forms that can modify the external environment in response to abiotic and biotic stimuli. Some exudate components contribute to soil adhesion (Akhtar et al., 2018; Galloway et al., 2018). Root epidermal and hair cells secrete compounds that contribute to root exudate profiles that can be plant species specific (Naveed et al., 2017). Because mutants in both root hair development and vesicle trafficking pathways showed root-substrate adhesion phenotypes (**Figure 2 and 3**), we asked if the centrifuge assay could identify effects of exudate composition on root-substrate adhesion. We selected Arabidopsis mutants reported to have altered exudate composition compared to wild type, including *pft1-3*, *jin1-9*, and *pdr2* (Badri et al., 2009; Berger et al., 1996; Carvalhais et al., 2015; Kidd et al., 2009). *PFT1* (*MED25*) encodes the MEDIATOR25 subunit of the Mediator nuclear protein, and *JIN1* (*MYC2*) encodes a basic helix-loop-helix leucine zipper transcription factor; both proteins are involved in the jasmonate signalling pathway (Kidd et al., 2009; Lorenzo et al., 2004). *pft1-3* and *jin1-9* mutants are reported to have altered root exudate composition, including lower amounts of the amino acids asparagine, ornithine and tryptophan than wild-type plants (Carvalhais et al., 2015). *PDR2* (*ABCG30*) encodes a pleiotropic drug resistance (PDR) full-length ABC transporter that is involved in ABA transport and the exudation of secondary metabolites (Badri et al., 2008; Kang et al., 2015). We did not observe root hair growth phenotypes in these mutant lines compared to wild type (**Figure 4A**).

**FIGURE 4.**
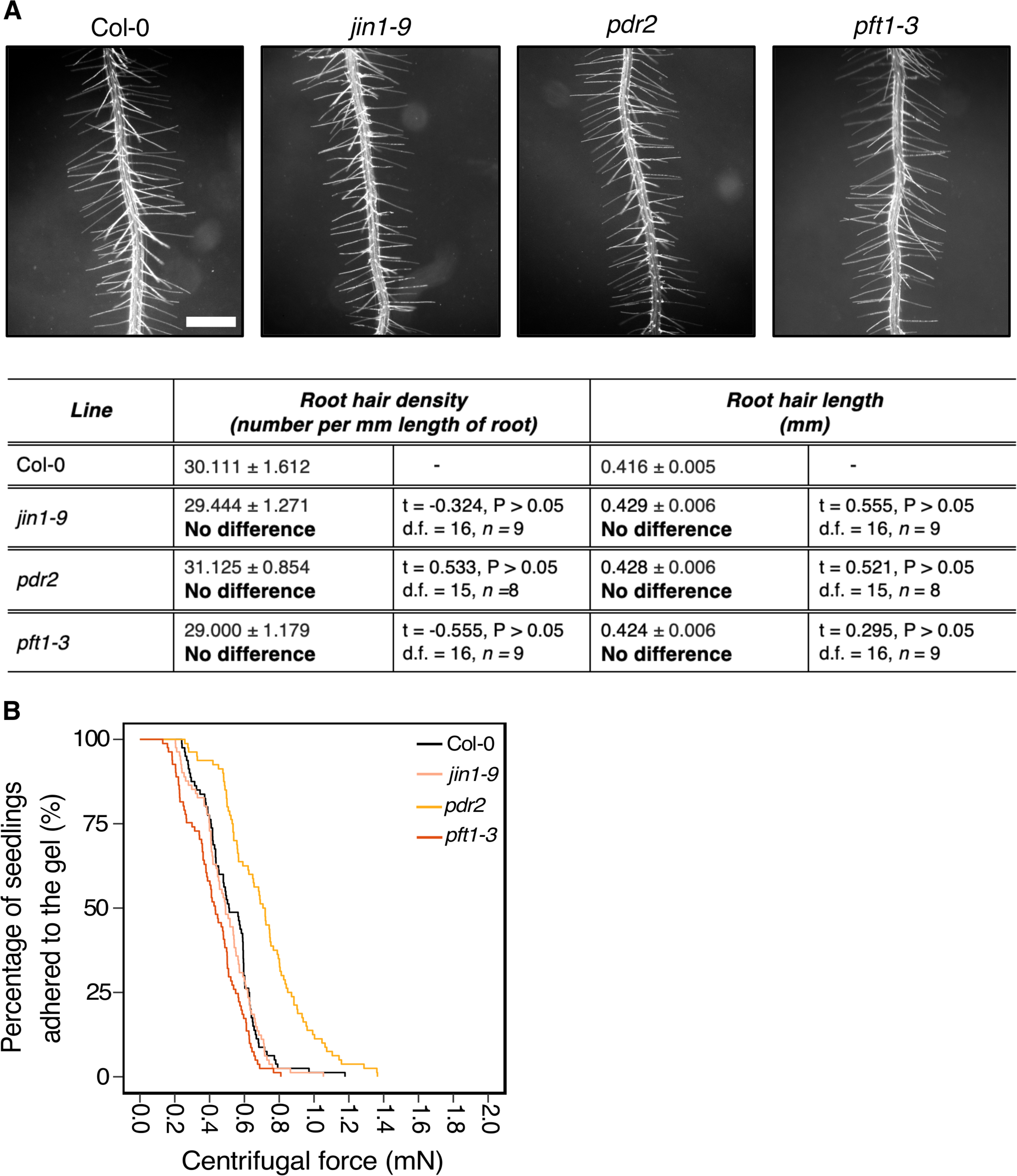
Exudate composition changes root-substrate adhesion properties in *Arabidopsis thaliana.* **(A)** Root hair phenotypes of 5-d-old wild-type (Col-0), *jin1-9, pdr2* and *pft1-3* seedlings grown on a gel medium and statistical comparisons of mean root hair density (number per mm length of root) and mean root hair length (mm) for each mutant line relative to wild type. Scale bar = 0.5 mm. In the table, the mean *±* standard error is given for the root hair density and root hair length of wild type and each mutant as well as a mean comparison to wild type, which is listed in bold. “No difference” is stated when there was no statistically significant difference between wild type and a mutant line. The statistical output of each univariate linear model is given. Alpha = 0.025. The results shown are from a representative experiment for at least two independent experiments. **(B)** Survival curves displaying the proportion of seedlings that adhered to the gel at increasing centrifugal force for 80 wild type (Col-0 – black), 81 *jin1-9* (light pink), 80 *pdr2* (light orange), and 81 *pft1-3* (dark orange). The results shown are from a representative experiment for at least two independent experiments showing a statistically significant difference in adhesion between mutant lines relative to wild type (Cox PH regression; alpha = 0.01), except for *jin1-9*. A single experiment included > 70 biological replicates for each candidate line.

Compared to wild-type seedlings, *pdr2* seedlings resisted detachment, with 0.38 times the risk of detaching from the gel (P<0.001 – **Table 2**; **Figure 4B**). Conversely, *pft1-3* seedlings had an increased risk of detachment 1.7 times that of wild-type plants (P < 0.001) and there was no difference in gel adhesion between *jin1-9* and wild-type plants (P>0.05) (**Table 2; Figure 4B**). These results show that exudate composition can alter root-substrate interactions, although this may not be the case for all exudate compositional changes, which may lay outside the sensitivity range of the assay.

### Centrifuge Assay as a Method for Forward Genetic Screening

Having demonstrated that our centrifugation technique can identify different types of traits that affect root-substrate adhesion, we used it in a forward genetic screen to find novel genes involved in root-substrate adhesion. To conduct this screen, the root-gel adhesion properties of individual plants from a pooled SALK T-DNA insertion mutant collection were analysed (Alonso et al., 2003). Individual plants that detached at the lowest centrifugal speed or remained attached to the gel after being spun at the highest centrifugal speed were recovered from the screen and self-fertilized to obtain progeny and identify the putative mutations. The centrifuge assay was used to rescreen the progeny (using a sample size of ≥ 70 *n*) to confirm whether candidate lines had significantly increased or decreased root-gel adhesion relative to wild-type plants. From this screen, we identified a line with significantly enhanced root-gel adhesion and conducted genomic Next-Generation sequencing followed by Sanger sequencing of T-DNA flanking PCR products to confirm a T-DNA insertion in the ABC transporter gene, *ABCG43,* whose function is unknown. We named this T-DNA insertion line *abcg43-1* and identified two additional mutant alleles, *abcg43-2* (SALK_201207c) and *abcg43-3* (SALKseq_30713) (**Figure 5A**).

**FIGURE 5.**
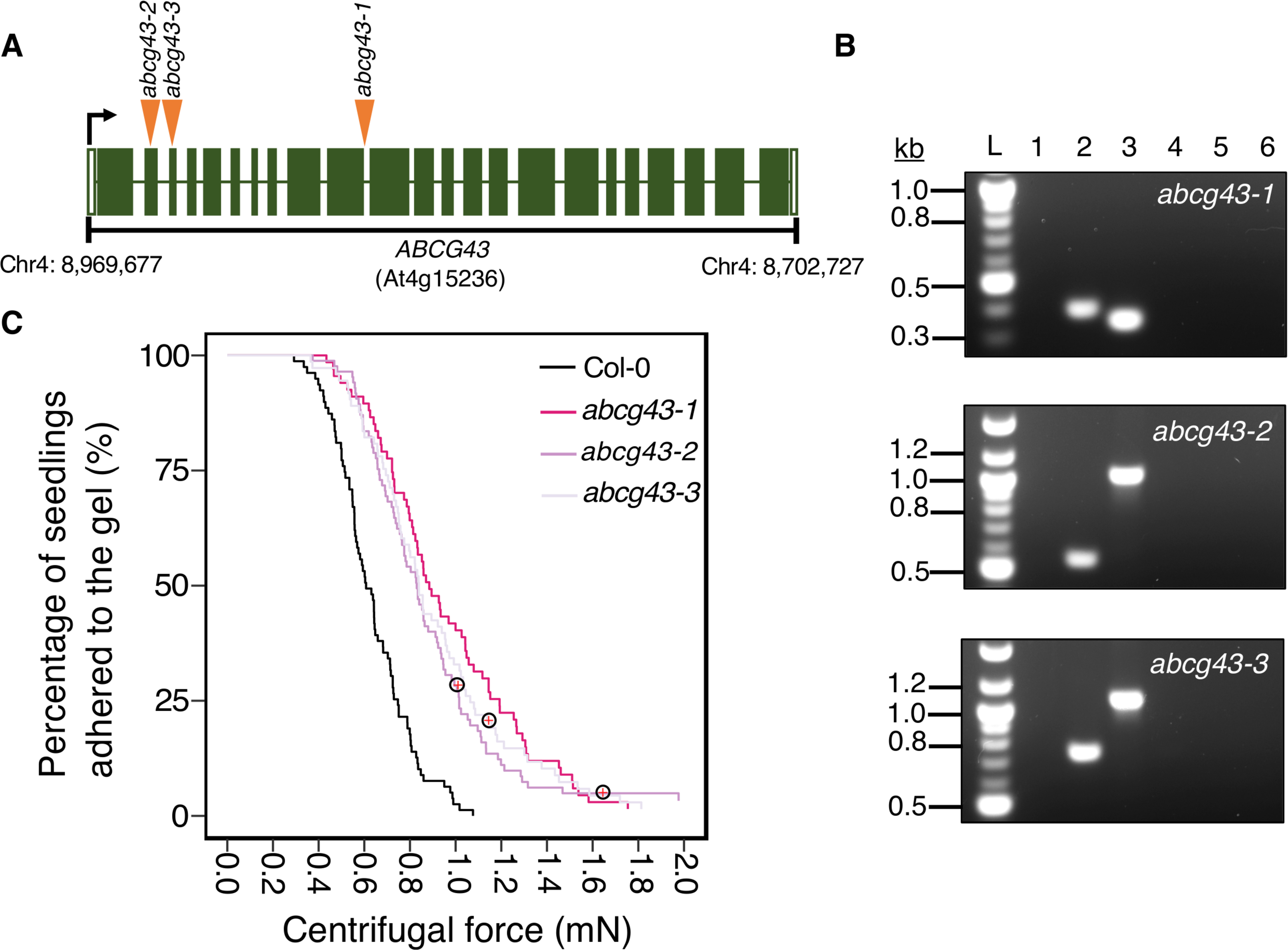
Using the centrifuge assay in a forward genetic screen to identify *Arabidopsis thaliana* root-substrate adhesion mutants. **(A)** T-DNA insert locations in *ABCG43* for each *abcg43* mutant allele, with insertions indicated by orange arrowheads. The *ABCG43* gene is located on chromosome four at position Chr4: 8,969,677-8,702,727 and contains 23 exons (dark green) and 22 introns (white gaps). *abcg43-1* has a T-DNA insert in the exon10/intron10 boundary, *abcg43-2* (SALK_201207) has a T-DNA insert in exon 2 and *abcg43-3* (SALKseq_30713) has a T-DNA insert in exon 3. **(B)** Genomic PCR confirming the homozygosity of each *abcg43* mutant for the respective T-DNA inserts. Lanes **1** and **2** were loaded with the gene-specific and T-DNA-border PCR products from *abcg43* mutant lines. Lanes **3** and **4** were loaded with the gene-specific and T-DNA-border PCR products from a wild type (Col-0) genomic DNA template. Lanes **5** and **6** were loaded with the water controls for the gene-specific and T-DNA-border PCR reactions, respectively. ‘**L’** indicates the 100-bp ladder. The expected product sizes for the gene-specific PCRs were ~350 nt, ~1025 nt and ~1090 nt; the T-DNA-border PCR product sizes were ~400 nt, ~550 nt and ~750 nt for *abcg43-1, abcg43-2* and *abcg43-3* alleles, respectively. **(C)** Survival curves displaying the proportion of seedlings that adhered to the gel at increasing centrifugal force for 79 wild type (Col-0 – black), 70 *abcg43-1* (dark pink), 85 *abcg43-2* (purple), and 73 *abcg43-3* (light purple). Red crosses circled on the survival curves represent seedlings that remained adhered to the gel after the maximum centrifugal speed (1611 RPM). The results are from a representative experiment for at least two independent experiments showing a statistically significant difference in adhesion between the mutant lines relative to wild type (Cox PH regression; alpha = 0.01). A single experiment included > 70 biological replicates for each candidate line.

We analysed the root-gel adhesion of the homozygous insertional *ABCG43* mutant alleles and found *abcg43-1*, *abcg43-2* and *abcg43-3* had detachment risks 0.25, 0.36 and 0.35 times that of wild-type plants, respectively (for all lines P < 0.001- **Table 2**; **Figure 5B, C**). These results illustrate the use of the centrifuge assay as a powerful screening method for identifying novel genes involved in root-substrate adhesion, as well as a tool for evaluating the effects of known gene function on root-substrate interactions.

## DISCUSSION

To date, most published methods that quantify the contribution of particular traits to plant-soil interactions are time-intensive and expensive (Bailey et al., 2002; De Baets and Poesen, 2010; De Baets et al., 2020; Toukura et al., 2006). In contrast, our assay can produce data within a week, is available to any laboratory with access to a bench top centrifuge and does not require speciality consumables. This method is a high-throughput and quantitative way to test the effects of root morphology and cell function on root-substrate adhesion and to screen for new biological and molecular factors that can alter plant-substrate interactions. Using a defined model system like Arabidopsis and sterile medium conditions permits the identification of plant-specific characteristics that can then be probed in other substrate- or soil-based conditions.

### Using the Centrifuge Assay to Investigate the Role of Root Hairs in Root-Substrate Interactions

Results presented in this paper show that the centrifuge assay can quantify effects of root hair morphology on plant-substrate adhesion. Previous work has shown that the presence of root hairs can significantly enhance root-substrate adhesion compared to the absence of root hairs (Choi and Cho, 2019; De Baets et al., 2020; Haling et al., 2013). We now show that our centrifuge assay can distinguish the adherence strength of seedlings with altered root hair shapes and sizes (**Figure 2**) and can quantify the contribution these morphological traits make to plant-substrate interactions. Some mutants with defective root hairs have mutations in cell wall biosynthesis and modification proteins, so it is possible that altered root-substrate interface compromises adhesion in these mutants, not just root hair shape. Using our method to rapidly phenotype mutants with root hair growth defects could direct additional experiments to characterize biochemical and morphological properties of genes of interest. Indeed, current evidence for the importance of root hairs for the deposition of complex polysaccharides with soil-binding properties (Galloway et al., 2020) supports the need for assays such as ours to expedite the identification of genes and mechanisms that improve soil-remediation, top soil maintenance, and erosion tolerance of at-risk land.

### Identifying New Genes Involved in Root-Substrate Adhesion

Mutations in cellular functions that do not alter root hair growth but do affect substrate adhesion can also be investigated using this assay. Vesicle trafficking pathways are required for cell wall deposition and maintenance (De Caroli et al., 2011; Larson et al., 2014; Rodriguez-Furlán et al., 2016); therefore, we hypothesized that mutants in vesicle trafficking pathways, particularly those at the plasma membrane, could affect root-substrate adhesion. Similarly, the composition and deposition of plant exudates help plants modify and optimize their growth conditions (Carvalhais et al., 2015; Naveed et al., 2017), suggesting that modification to plant exudate composition could affect root-substrate interactions. Consistent with these hypotheses, mutants with vesicle trafficking defects and altered exudate composition had root-substrate adhesion properties that differed from wild type, indicating that this method can identify aspects of plant cell biology important for root-substrate interactions that do not affect root or root hair morphology (**Figures 3 and 4**). Our results further indicate that while root hairs provide significant adhesive and cohesive support both on solid gel and soil media (De Baets et al., 2020), there are additional cellular factors contributing to plant-substrate interactions that are not directly observable without the use of expensive reagents, development of time-intensive transgenic lines, and microscopy equipment. Our centrifuge assay provides an affordable and quick alternative or complement to these current methodologies.

The compounds and interactions that contribute to root-substrate adhesion are complex, and the direct and indirect factors that participate in these interactions still remain to be characterised in detail. Many root cell wall or secreted compounds might directly affect adhesive characteristics of the root surface, such as arabinogalactan proteins involved in the adhesion of English ivy (Huang et al., 2016). Other complex polysaccharides have also been shown to enhance soil binding in both Arabidopsis and cereal root systems (Galloway et al., 2018, 2020). Indeed, mutants in genes associated with the production of these compounds would be potentially excellent candidates for the application of this assay, which would quantify the contribution these compounds make to root-substrate adhesion.

We cannot expect all exudate components to present adhesive phenotypes because not all will affect root-substrate adhesion, and some will affect adhesion in a way that is not detected by this assay. For example, the *jin1-9* mutant tested in our exudate cohort was not significantly more or less adhesive compared to wild-type roots (**Figure 4**) despite its altered exudate composition (Anderson et al., 2004). Some exudate components will play no role in adhesion, while others may require additional co-factors present in soil or non-sterile media to participate in root-substrate interactions. Although these are examples of the limitations of the assay, this assay does allow for the identification of direct contributions that plants make to root-substrate cohesion and its robust statistical power and high-throughput aspects will significantly improve genetic screening timelines and enhance large-scale experimental design.

Forward genetic screening relies on distinctive phenotypes that distinguish differences within large, genetically diverse populations. High throughput, creative techniques for identifying phenotypes within morphologically similar populations, such as thermal imaging (Plessis et al., 2011) and confocal imaging platforms (Drakakaki et al., 2011; Robert et al., 2008) require specific equipment that not all laboratories have at their disposal. In contrast, the centrifuge assay can be used as a screen to identify new genes affecting plant-substrate interactions without specialist equipment, making it an accessible option for research groups that may not otherwise be able to efficiently conduct genetic screens. Moreover, the centrifuge assay does not require morphological or conditional differences for candidate selection. As an example, we report the identification in a forward genetics screen of a mutant population in which *abcg43* mutant seedlings were identified and shown to be more adhesive to gel than wild type (**Figure 5**). The function of ABCG43 in Arabidopsis is uncharacterised, indicating that this assay can identify novel candidate genes that have no previously known association with root morphological defects, adhesion or exudate composition for future study.

## CONCLUSION

This assay has applications in plant cell biology and genetics to investigate how diverse gene activities, cell functions and root structure affect plant-soil adhesion, and identify the molecular pathways involved. Potential applications not tested in this report include the measurement of the effects environmental conditions have on plant-substrate adhesion and how other plant species or ecotypes respond to detachment forces. Given the recent discovery that root hair presence is important for Arabidopsis cohesion between roots and gel, and compost or soil (De Baets *et al.,* 2020), this assay can broadly contribute to plant and soil sciences through the identification of plant traits that increase root-substrate cohesion and potentially stabilise slopes or reduce soil erosion.

## SUPPLEMENTARY INFORMATION

**Table S1** Centrifuge assay troubleshooting

**Table S2** Primers used in this study for genotyping *Arabidopsis thaliana* mutants

## ACKNOWLEDGEMENTS

This work was supported by a Leverhulme Trust project grant RPG-2013-260 to CSG and TBL. BME was supported by a BBSRC SWBio PhD studentship (grant BB/M009122/1). We are thankful to Don Grierson FRS and Enrico Coen FRS who independently suggested the centrifuge assay concept, Timothy Quine for providing insightful guidance during method development, and Thomas Denbigh for providing technical advice. We are grateful to Jill Harrison, Victoria Spencer, Zoe Nemec Venza, Sophie Carpenter and Nicholas H Fair for their helpful comments during revision of the manuscript. This manuscript has been released as a pre-print at [http://doi.org/10.1101/2020.05.14.095448] (Eldridge et al., 2020).

## AUTHOR CONTRIBUTIONS

CSG designed the initial project, conducted proof-of-concept pilot experiments, obtained funding, and recruited and advised the team. TBL and IC developed the theoretical parameters for the centrifugal assay and the bounds on the errors from other mechanical forces. BME and ERL planned and conducted all experiments apart from the initial genetic screen, which was carried out by LW and KMS. BME and ERL analysed all of the data presented herein. AS and BME optimised the centrifuge assay data collection and data analysis methods. BME and ERL wrote the initial manuscript. BME, ERL, and CSG edited the manuscript. All authors reviewed and approved the final manuscript.

