## Supplemental Tables 1 and 2 for "A centrifuge-based method for identifying novel genetic traits that affect root-substrate adhesion in *Arabidopsis thaliana*" (from another version of this preprint: version 4, 2020-10-29)

### *Supplementary Material*

**Table S1** Centrifuge assay troubleshooting

| ***Problem observed*** | ***Possible reason(s)*** | ***Possible solution(s)*** |
|---|---|---|
| Candidate lines and wild-type seedlings significantly differ in size (e.g. in cotyledon size and root length). | Differences in germination rates and development rates of particular lines (e.g. due to age of seed stocks). It is important for candidate lines and wild-type plants to be comparable in size and at the same developmental stage for comparisons of gel-adhesion to be valid. | 1. Set up a time-lapse experiment to compare the developmental stage of candidate line(s) to wild-type seedlings.<br>2. Decrease or increase the stratification periods of particular lines.<br>3. Extend the growing period of particular lines (by e.g. +12 hrs for lines developing slowly or -12 hrs for lines developing quickly).<br>4. Generate new seed stocks. |
| The seedlings are growing into the gel. | The gel has been pierced when the seed was sown or the angle at which the Petri plates have been orientated during the growing period was above or below ~80° causing the roots to grow into the gel. | Regularly check that seedlings grow down the surface of the gel and correct the plate angle if necessary. |
| The gel is shattering during centrifugation | Possible causes include:<br>1. The medium was too hot when poured so water evaporated, resulting in a higher than intended and/or uneven gel strength.<br>2. The medium was reheated prior to pouring which changed the gel composition.<br>3. The surface of the gel was pierced during seed planting, initiating local gel fractures.<br>4. The Petri plates have been roughly handled, causing fractures.<br>5. The acceleration and/or deceleration speed on the centrifuge was too high. | 1. Pour the medium into the Petri plates when the media has cooled significantly (and is not steaming). Ensure the medium is thoroughly mixed and bubble free when poured.<br>2. Do not reheat medium. Store in an oven at 50-60°C to prevent solidification prior to pouring.<br>3. Use a desk lamp and magnifier when sowing plates to ensure you do not pierce the surface of the gel.<br>4. Discard plates that have been dropped (even though the gel may not have visibly shattered).<br>5. Prior to conducting experiments, check the rate of gel shattering (using plates that have not been sown with seed) at different acceleration/ deceleration speeds. |

**Table S1** continued…

| | | |
|---|---|---|
| Replicate size is too low. | 1. Poor germination/ seedling growth.<br>2. Insufficient replicate Petri plates to ensure an adequate replicate size. | 1. See above for issues relating to germination/ seedling growth.<br>2. Sow spare Petri plates (e.g. we sow 150 seeds across 15 Petri plates per line to guarantee >70 biological replicates for each line in each experiment). |
| The adhesion properties of lines have changed. | Because roots are extremely sensitive to their environment, wild-type adhesion properties may shift between experiments.<br>1. Gel batches differ or environmental conditions including light, temperature or water quality fluctuated.<br>2. Pouring growth media too hot or incubating plates in unsuitable growth spaces can cause liquid to accumulate on the surface of the gel, disrupting adhesion,<br>3. A different wild-type stock was used without checking the adhesion properties of the stock. | 1. Void any experiments where excess surface water has accumulated on plates.<br>2. Check the growth conditions are consistent (e.g. batches of growth medium, water quality, light and temperature within the growth cabinet, etc.).<br>3. Ensure you run technical repeats of the assay to confirm the adhesion properties of a candidate line relative to wild-type plants are consistent.<br>4. Use different pooled wild-type seed stocks for a technical repeat of an experiment to ensure that the adhesion phenotype of a candidate line is consistent when a different wild-type stock is used.<br>5. Check and compare the adhesion properties of wild-type stocks in a single preliminary experiment. |

**Table S2** Primers used in this study for genotyping *Arabidopsis thaliana* mutants

| ***Oligo name*** | ***Oligonucleotide sequence (5' to 3')*** | ***Allele*** |
|---|---|---|
| *jin1-9_F* | GGCGGGATTTAATCAAGAGAC | SALK_017005 |
| *jin1-9_R* | TTTGGTACAACCGCTCGTAAC | |
| *pft1-3_F* | CGATCGAGTTGACCAAAGAAG | SALK_059316 |
| *pft1-3_R* | TTTGCATCAGGCAATATGTTG | |
| *pdr2_F* | ATGATCCAAACAGGTGAAGAAG | SAIL_F08_811 |
| *pdr2_R* | CTATTTCTTTTGGAAACTGAGTTTGC | |
| *abcg43-1_F* | GGGCTACAACTACAAGACGAAC | N75206[b] |
| *abcg43-1_R* | GGGAAAAGAACAAAGAACCCAAAAG | |
| *abcg43-2_F* | CTCGTGAAGCCAACTTGCTAG | SALK_201207 |
| *abcg43-2_R* | TCTGTAGAGTGGAAGCAACCC | |
| *abcg43-3_F* | CCCTCAATAGAAGCTGCCTG | SALKseq_30713 |
| *abcg43-3_R* | TGTTGAGGATGATAATCGCG | |
| SAIL_LB3 _B | TAGCATCTGAATTTCATAACCAATCTCGATACAC | SAIL_F08_811 |
| SALK_LBb1.3_B | ATTTTGCCGATTTCGGAAC | N75206[a] SALK201207<br>SALKseq_30713<br>SALK_017005<br>SALK_059316 |

'_F' or '_R' refer to the forward and reverse gene-specific primers and '_B' refers to the T-DNA-specific border primer. [a]The *abcg43-1* line was identified from a pooled set of 100 SALK T-DNA insertion lines (stock number N75206, Alonso *et al.,* 2003).
